# Neural characterization of the “totonou” state associated with sauna use

**DOI:** 10.1101/2023.01.06.522959

**Authors:** Ming Chang, Takuya Ibaraki, Yasushi Naruse, Yasuhiko Imamura

## Abstract

Saunas are becoming increasingly popular worldwide, being an activity that promotes relaxation and health. Intense feelings of happiness have been reported shortly after enjoying a hot sauna and cold water, what is known in Japan as the “totonou” state. However, no research has investigated what occurs in the brain during the “totonou” state. In the present study, participants underwent a sauna phase, consisting of three sets of alternating hot sauna, cold water, and rest. We elucidated changes in brain activity and mood in the “totonou” state by measuring and comparing brain activity and emotional scales before and after the sauna phase and during the rest phase in each set. We found significant increases in theta and alpha power during rest and after the sauna phase compared to before the sauna phase. Moreover, in an auditory oddball task, the p300 amplitude decreased significantly and MMN amplitude increased significantly after the sauna phase. The increase in MMN indicates higher activation of the pre-attentional auditory process, leading to a decrease in attention-related brain activity P300. Hence, the brain reaches in a more efficient state. Further, the response time in behavioral tasks decreased significantly. In addition, the participants’ subjective responses to the questionnaire showed significant changes in physical relaxation and other indicators after being in the sauna. Finally, we developed an artificial intelligence classifier, obtaining an average accuracy of brain state classification of 88.34%. The results have potential for future application.

## Introduction

Sauna is a type of steam and hot air bath, possibly originated in Finland, where there are builtin saunas in almost every house [1]. The indoor temperature of the sauna room ranges from about 70°C to over 110°C. After sauna bathing, it is typical to cool down by immersion in the snow or water or by taking a break in the outside, before re-entering the sauna. Essentially, sauna therapy takes advantage of the thermoregulatory properties of mammals and birds [2]. Regular sauna use may cause profound physiological effects and could potentially stave off neurodegenerative diseases by different mechanisms. In the Kuopio Ischemic Heart Disease (KIHD) Risk Factors Study, a 20.7-year ongoing, population-based prospective cohort health study conducted among >2300 middle-aged men from eastern Finland [3] found an association between sauna use and a reduced risk of age-related diseases, including cardiovascular disease, neurodegenerative disease, metabolic dysfunction, and decreased immune function. The study also demonstrated that men who used the sauna 4–7 times per week had a 65% reduced risk of developing Alzheimer’s disease than those who used the sauna only once per week. Sauna use has been shown to reduce symptoms of depression [4] and promote transient growth hormone release, an effect which varies with time, temperature, and frequency of exposure [5]. Sauna use promotes a strong increase in β-endorphins [6,7], which seem to be partly responsible for the euphoria associated with exercise [8]. Moreover, sauna use may reduce the risk of certain chronic or acute respiratory diseases, including pneumonia [9]. As well as reduce the incidence of the common cold and improve respiratory health [10,11]. Other findings point to the effects of sauna use on the immune system and heat shock proteins. A single Finnish sauna session increased white blood cell, lymphocyte, neutrophil, and basophil counts in athletes and non-athletes; but to a higher degree in athletes. [12]. The heat stress from saunas can improve physical health by increasing cardiopulmonary fitness and endurance and maintaining muscle mass [13].

However, there is little research on saunas from the perspective of neuroscience and psychology. A recent study examined the effects of saunas on brain activity and found that post-sauna recovery enhances brain relaxation and improves cognitive efficiency in oddball tasks [14]. However, since changes in participants’ subjective states (mood, etc.) were not examined, it is unclear whether changes in brain activity are associated with the improved mood. In addition, as only brain activity was measured before and after the sauna, it is not clear what occurs during the time in the sauna.

People feel happy after being in the sauna; in fact, some individuals report a strong, but short sense of well-being after being in the sauna, known as a “totonou” state in Japan. The word “totonou” means to be prepared /arranged in Japanese; with respect to the sauna, the “totonou” is the well-being state in which the body and mind are automatically conditioned by entering the sauna, and the original ability of the person is restored. To achieve the “totonou” state, the body needs to be repeatedly heated and cooled, with breaks in-between. In general, it is important to perform ≥3 sets of alternating hot and cold bath including the following three stages: sauna (hot) → water bath (cold) → outdoor rest’ for reaching the “totonou” state, which usually occurs at the rest step. What exactly happens in the “totonou” state in the brain remains unclear. Thus, both qualitative and quantitative brain and behavioral data in the “totonou” state are required. The components of event-related potentials (ERPs) observed using the oddball paradigm reflect sensory and cognitive processes in brain activity [15]. In the oddball paradigm, two different stimuli are presented in various ways; the subject is instructed to respond to the infrequent or target stimulus. As for the ERP components, we focused on P300 and mismatch negativity (MMN) during the auditory oddball paradigm. P300 is one of the positive components of the ERP and typically appears at 250–400 ms following target stimulus presentation. The P300 amplitude is influenced by attention such that greater attention produces a larger P300 [16,17,18]. Moreover, MMN appears in the auditory oddball paradigm at 100–250 ms following target stimulus presentation [19,20]. Unlike P300, MMN can be elicited without the listener subjectively attending to the sound stimuli [21-23]; therefore, the MMN is known to be caused by the neural activity of a pre-attentive auditory process [24,25].

This study aimed to verify the effect of sauna bathing through an objective brain activity and subjective mood evaluation. In particular, we aimed to elucidate changes in brain activity and mood in the “totonou” state by measuring and comparing brain activity and mood-related mind assessment scales before and after entering the sauna, and at rest between each set. In addition, we developed an artificial intelligence (AI) classifier based on electroencephalography (EEG) and verified its accuracy in classifying brain states (totonou or non-totonou).

## Methods

### Participants

Twenty healthy participants were recruited for participation in the experiment (14 men; 21–41 years old). All participants were right-handed. None of the participants had hearing or speech disorders.

The participants were assigned to two groups: a sauna group and a control group. The sauna group consisted of eight men and two women, and the control group consisted of six men and four women. All subjects in the sauna group had experienced sauna bathing. Informed consent was obtained from all participants prior to the study. All experimental procedures were approved by the Shiba Palace Clinic Ethics Review Committee. Additionally, all procedures were in accordance with the Helsinki Declaration of 1964 and later revisions.

### Procedure

The experimental procedure in the sauna group consisted of a pre-sauna phase, sauna phase, and a post-test phase (Fig. 1). Three sets of 3 steps alternating hot and cold baths: sauna (hot, 85∼90°C) → water bath (cold, 16°C) → rest (20°C) was performed during the sauna phase. One set consisted of sauna (10 min), water bath (1∼2 min) and rest (7 min). During the rest stage, the participants were asked to close their eyes and relax. We measured scalp EEG and heart rate (HR) during an auditory oddball task in the pre-sauna and post-sauna phases. Similarly, we also measured salivary a-amylase activity (SAA) to evaluate participants’ stress using an SAA biosensor (salivary amylase monitor, NIPRO Co., Japan) in the pre-sauna and post-sauna phases [26]. In addition, in-ear EEG was performed for frequency analysis and the responses to questionnaires related to physical and mental states were recorded in the second half of the pre-sauna phase and rest immediately after the cold water bath during the sauna phase.

**Fig. 1.**
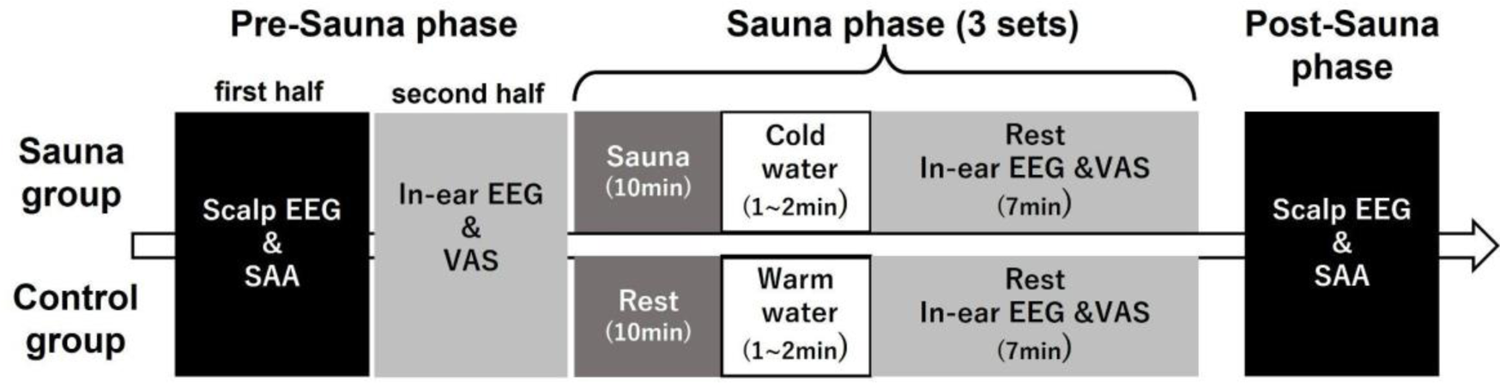
Experimental procedure

In the control group, the participants did not take saunas but rested. The participants in the control group entered a warm water bath (37°C) during the bath phase. This was designed to allow participants in both groups to have the same exposure time of skin to water.

### Oddball Paradigms

In the auditory oddball task, a pseudorandom series of frequent standard stimuli (1000 Hz tone; 80%) and infrequent “target” (2000 Hz tone; 20%) stimuli, were presented with a 1.5-s stimulus onset asynchrony (SOA). The auditory stimuli were 50 ms (5 ms rise/fall time) and presented via headphones at a sound pressure level of 60 dB. The serial order of the stimuli was pseudo-random with the restriction that ≥2 standard stimuli were presented between infrequent “target” stimuli. During the auditory oddball tasks, participants were instructed to press a response button to target stimuli with their preferred hand. Reaction times (RTs, in ms) were measured and only correct responses were averaged. A total of 200 stimuli were presented in the task.

### Electroencephalographic recording

We need to immediately record the EEG in rest after the cold water bath during the sauna phase, which is not possible with conventional EEG record devices as a short span of time is available to install the EEG devices. Hence, two types of EEG record device were used in this experiment. EEG was recorded using a Wireless Biosignal Amplifier System (Polymate Mini AP108; Miyuki Giken Co., Ltd., Tokyo, Japan) and solid gel electrodes (METS INC., Chiba, Japan) during the oddball paradigm conducted in the pre-sauna and post-sauna phases. Hereafter, this EEG data will be referred to as scalp EEG data. It was recorded (500 Hz sampling rate) from Cz, Fz and Pz, according to the international 10–20 system. In addition, ground and reference electrodes were placed on the left and right mastoids, respectively. Electro-ocular (EOG) activity was assessed using one electrode placed at the upper-outer edge of the left eye to measure eye blinks and vertical eye movements. An electrocardiogram (ECG) was obtained using one electrode placed at the left upper chest to measure HR. Data recorded during the oddball task were filtered using 0.1 Hz low cutoff and 30 Hz high cutoff filters. The epoch for the standard or target stimulus started 100 ms before stimulus presentation (baseline) and continued 600 ms after stimulus onset. Epochs in which the EEG or EOG signal > ±50 μV due to vertical eye movements and muscle contraction artifacts were rejected automatically; then, the standard and target ERP were obtained by averaging the epochs for each stimulus. The difference waveform was calculated by subtracting the standard ERP from the target ERP. We calculated the area of mismatch negativity (MMN) as the negative peak in the difference waveform at 100–250 ms and the area of P300 as the positive peak in the difference waveform at 240–400 ms from stimulus onset.

In addition, EEG was recorded from the left and right ear canals using an in-ear EEG System (VIE ZONE, VIE STYLE Inc., reference electrode: the back of the neck) as it has the advantage of a very short installation time. Hereafter, this EEG data will be referred to as In-ear EEG data [27]. The sampling rate of EEG data was 600 Hz. In-ear EEG data were filtered using 2 Hz low cutoff and 30 Hz high cutoff filters. Data were divided into equal sized segments (8 s); segments in which the EEG signal > ±100 μV due to vertical eye movements and muscle contraction artifacts were rejected automatically; then, Fourier transformations were conducted. To overcome the limitation of the individual specificity and incomparability in the traditional EEG frequency band, we used the individual alpha frequency (IAF)-based frequency band division method [28]. The peak power frequency among 8–13 Hz obtained from the data during the pre-sauna phase was defined as IAF. Frequency bands were determined individually for each subject using IAF as cutoff point between the lower and upper alpha bands. Each participant’s data were averaged across epochs for the left ear and right ear channels respectively, and the gross absolute spectral power was computed for five frequency bands:

1. theta band: [IAF − 4.0 Hz to IAF − 6.0 Hz];
2. lower 1 alpha band: [IAF − 2.0 Hz to IAF − 4.0 Hz];
3. lower 2 alpha band: [IAF to IAF − 2.0 Hz];
4. upper alpha band: [IAF to IAF + 2.0 Hz];
5. beta band: 15–30 Hz

### Questionnaires

In the second half of the pre-sauna phase and immediately after In-ear EEG measurement, the participants answered a questionnaire based on a visual analogue scale (VAS) on the PC. In the questionnaire, we asked participants to answer 14 questions extracted from the “Altered states of consciousness rating scale [29]” and a short-form self-report measure to assess relaxation effects (S-MARE) [30]. The problem list is shown in Tables 2 and 3. The S-MARE includes 15 items that evaluate psychological and physiological aspects and is composed of three subscales: “physiological tension,” “psychological calm,” and “cognitive tension”.

**Table 1.** Summary of post-hoc tests for pairwise comparisons of sets on each frequency band. * p < 0.05.

|  | Theta | Lower1 alpha | Lower2 alpha | Upper alpha | Bata |
| --- | --- | --- | --- | --- | --- |
| pre — post1 | * | * | n.s. | * | n.s. |
| pre — post2 | * | * | * | * | n.s. |
| pre — post3 | * | * | * | * | n.s. |
| post1 — post2 | n.s. | * | n.s. | n.s. | n.s. |
| post1 — post3 | n.s. | n.s. | n.s. | n.s. | n.s. |
| post2 — post3 | n.s. | n.s. | n.s. | n.s. | n.s. |

**Table 2.** Results of two-way repeated measures ANOVA for all questions in “Altered states of consciousness rating scale”. **+** p < 0.1, * p < 0.05 and ** p < 0.01.

| questionnaires |  | interaction (group × set) | group effects | set effects |
| --- | --- | --- | --- | --- |
| No, not more than usually | Yes, much more than usually | F value | F value | F value |
| • Sounds seemed to influence what I saw. |  | 1.76 | 0.02 | 4.38 ** |
| • I experienced boundless pleasure. |  | 6.66 ** | 0.38 | 16.37 ** |
| • Some everyday things acquired special meaning. |  | 4.27 * | 1.99 | 6.84 ** |
| • I could see images from my memory or imagination with extreme clarity. |  | 19.46 ** | 0.38 | 8.32 ** |
| • I felt as if I was floating. |  | 6.21 ** | 0.38 | 12.43 ** |
| • I saw brightness or flashes of light with closed eyes or in complete darkness. |  | 3.51 * | 0.02 | 4.06 * |
| • I felt one with my surroundings. |  | 1.28 | 6.87 * | 16.64 ** |
| • I felt isolated from everything and everyone. |  | 8.59 ** | 0.91 | 10.61 ** |
| • I had very original thoughts. |  | 1.38 | 1.06 | 6.46 ** |
| • Bodily sensations were very enjoyable. |  | 2.76 + | 0.12 | 13.68 ** |
| • my muscles feel loose. |  | 3.03 * | 0.14 | 2.42 + |
| • my breathing is faster than usual. |  | 11.44 ** | 2.06 | 6.69 ** |
| • I’m feeling very relaxed. |  | 4.44 ** | 0.11 | 14.45 ** |
| • In the “totonou” state |  | 6.54 ** | 0.64 | 28.21 ** |

### Statistical analysis

For the scalp EEG data, a repeated measures multivariate analysis of variance (ANOVA) was used to determine the effects of sauna bathing with electrode location (Fz vs. Cz vs. Pz), group (sauna vs. control), and stage (pre vs. post3) as factors influencing cognitive processing (characteristics of ERPs components).

For In-ear EEG data, a repeated measures multivariate ANOVA was used to determine the effects of sauna bathing with electrode location (L vs. R), group (sauna vs. control), and sauna set (pre vs. post1 vs. post2 vs. post3) as factors influencing brain neural oscillations (theta, low1 alpha, low2 alpha, upper alpha, and beta).

A two-way repeated measures ANOVA was used to assess the change of subjective feelings about their physical and mental state with group (sauna vs. control) and sauna set (pre vs. post1 vs. post2 vs. post3) as factors affecting the questionnaires’ score.

A two-way repeated measures ANOVA was used to assess the other effects of sauna bathing with group (sauna vs. control) and stage (pre vs. post3) as factors affecting cognitive performance (reaction time), SAA, and heart rate (HR).

If significant effects were found, the Holm–Bonferroni method was used as a post-hoc test for multiple comparisons within each repeated-measure ANOVA. The p-values for them was <0.05.

### AI decoding algorithm

In-ear EEG data were categorized into two categories: In-ear EEG data in the pre-sauna period were “non-totonou” and the other In-ear EEG data were “totonou”. This means that the “totonou” category data includes data collected during three rests in the sauna phase (post1, post2, and post3), while the “ non-totonou” category data only includes data collected during the pre-sauna phase (pre).

In-ear EEG data were divided into equal sized segments (4 s) and a Fourier transform was conducted. The power of the 4–40 Hz bands was calculated at 0.5 Hz intervals. Noise quantification was performed for each segment based on our pre-setting criteria, derived from clean In-ear EEG data [27]; segments > 2SD were excluded. Then, the power was standardized within participants. For future applications, three signals (left ear, right ear, and the difference between the signals of the left and right ears) were used for classification between “totonou” and “non-totonou” categories. Since the number of “totonou” category data was larger than that of “non-totonou” category, the data in the “totonou” category were randomly selected to include the same number of data as in the “non-totonou” category. The number of feature dimensions was reduced by applying principal component analysis (PCA) to all components from 1st to 150^th^ component. The training data included 90% of each category data and 10% of each category data were test data. Linear discriminant analysis (LDA) with least absolute shrinkage and selection operator (LASSO) was adopted for discrimination of the two categories [31]. A 10-fold cross-validation of training data was performed to select optimal parameter lambda (λ), the optimal value is considered to be the lambda value that returns the lowest testing error over the cross-validation, and finally the accuracy was evaluated using test data.

## Results

### Effects of sauna on cognitive processing

We assessed whether cognitive processing changed after sauna bathing. Using scalp EEG data collected during the pre-sauna and post-sauna (post3), we calculated and compared the P300 and MMN amplitudes for all participants in the sauna and control groups (Fig. 2).

**Fig. 2.**
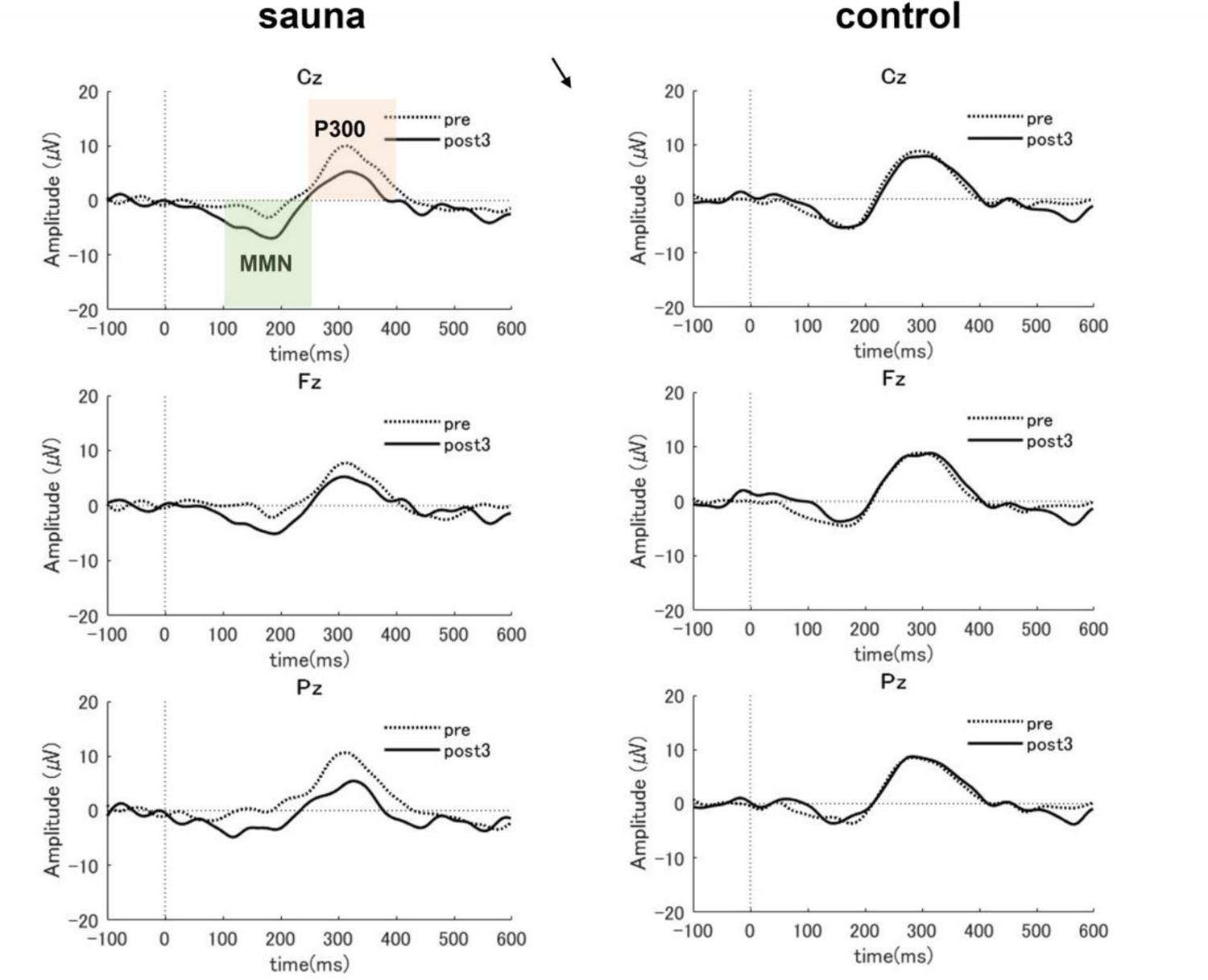
Grand-averaged event-related potential (ERP) difference waveform at three EEG sites (Fz, Cz, and Pz) for auditory tasks before and after the sauna in the sauna group vs. control group.

For the P300 area, the repeated measures ANOVA indicated a significant main effect of stage (F (1, 54) = 12.61, p < 0.01), a significant main effect of group (F (1, 54) = 5.16, p < 0.05) and a significant interaction between group and stage (F (1, 54) = 16.17, p < 0.01). We found no significant effects for electrode location or other interactions. We examined the simple main effects of group and stage to decompose the significant group × stage interaction. A simple main effects test for the stage showed that the P300 area significantly decreased in the sauna group (F (1, 54) = 28.68, p < 0.01) but not in the control group (F (1, 54) = 0.11, n.s.; Fig. 3). For the MMN area, the repeated measures ANOVA indicated a significant main effect of stage (F (1, 53) = 23.00, p < 0.01) and a significant interaction between group and stage (F (1, 54) = 26.50, p < 0.01). We found no significant effects for electrode location or other interactions. We further examined the simple main effects of group and stage to decompose the significant group × stage interaction. A simple main effects test for the stage showed that the MMN area significantly increased in the sauna group (F (1, 53) = 50.47, p < 0.01) but not in the control group (F (1, 53) = 0.03, n.s.; Fig. 3).

**Fig. 3.**
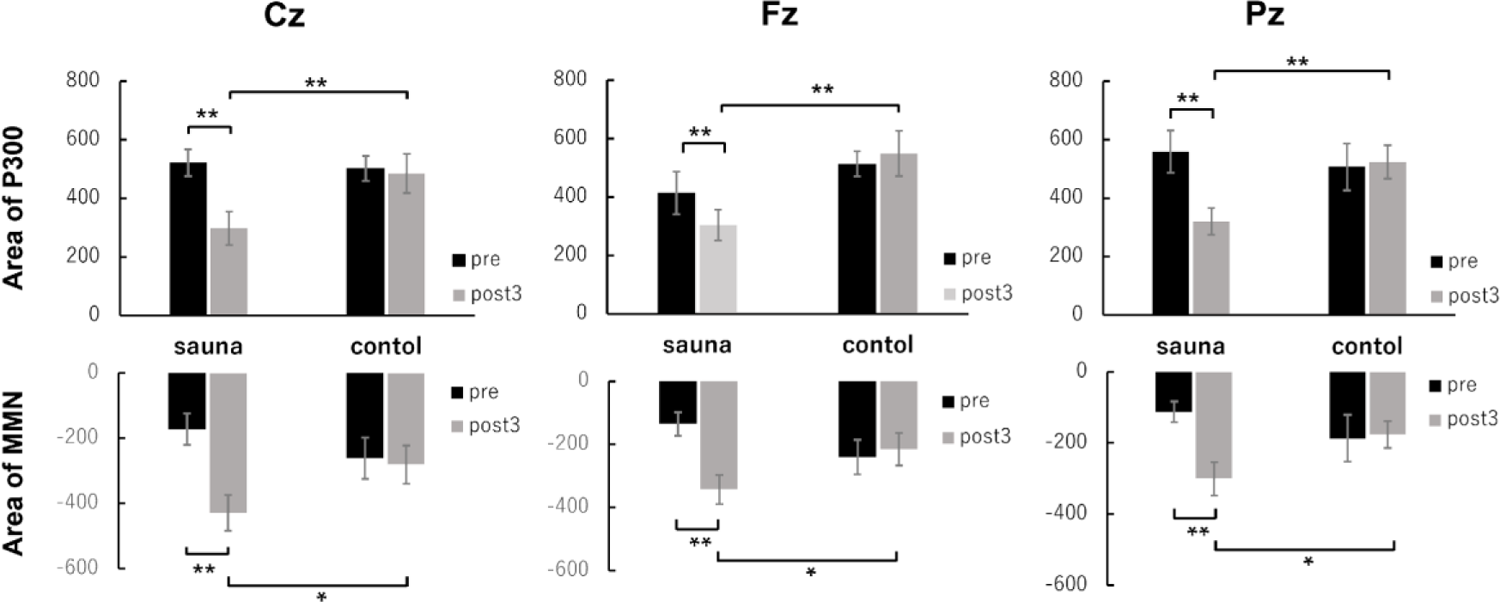
Average MMN and P3 amplitudes at three sites (Fz, Cz, Pz) before (pre) and after sauna (post3) in the sauna and control groups. Error bars represent s.e.m.* p < 0.05 and ** p < 0.01.

### Effects of sauna bathing on neural oscillation

We assessed whether neural oscillation changed during the sauna bathing set. Using ear EEG data collected during the pre-sauna phase and during each rest in the sauna phase, we calculated the amplitude spectral power in the sauna and control groups (Fig. 4). In addition, we calculated and compared the gross absolute spectral power of five frequency bands in the sauna and control groups.

**Fig. 4.**
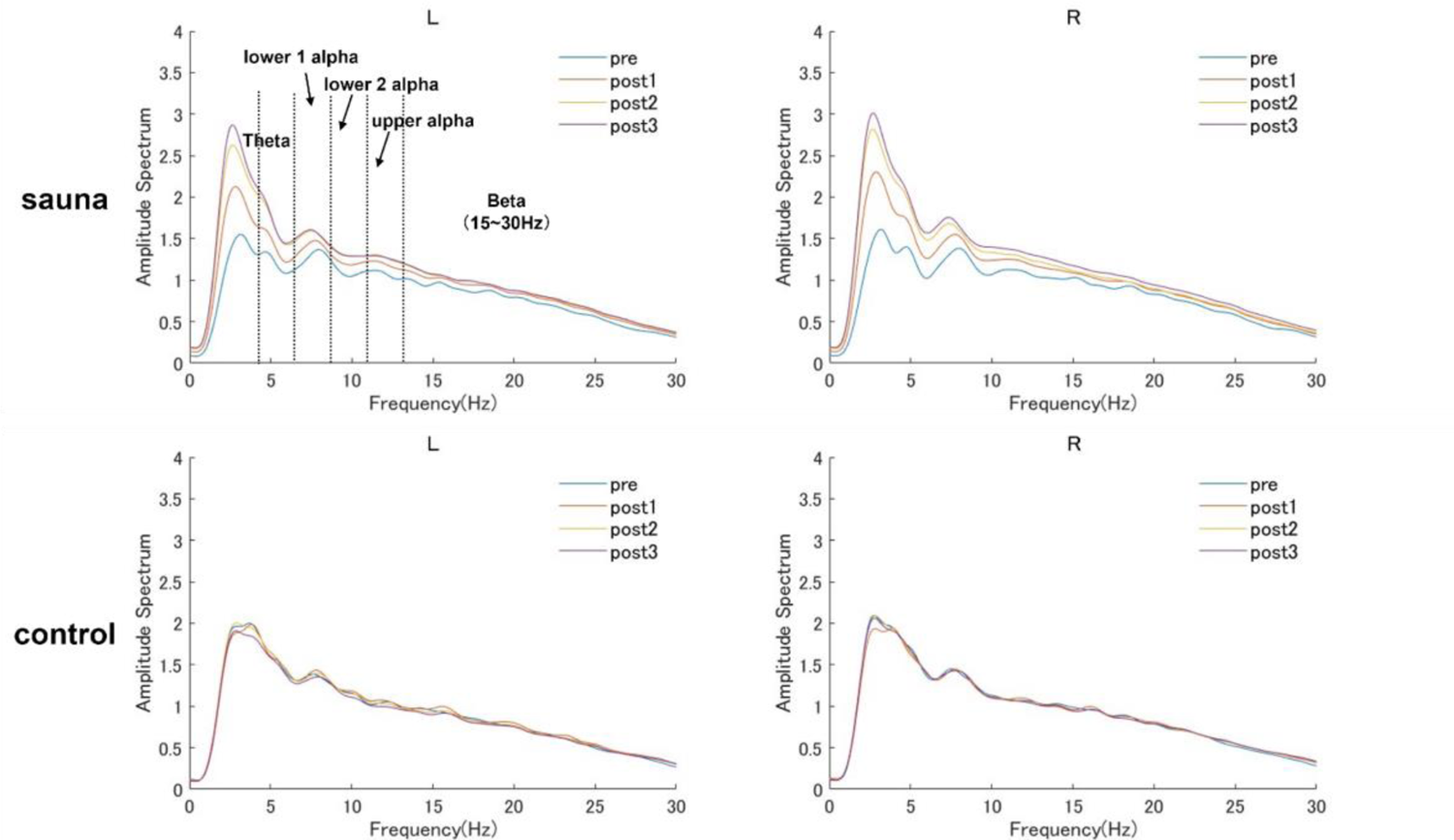
Amplitude spectral power for the L and R electrodes (top) in the pre-sauna (pre) and at each rest in the sauna set (post1, post2, post3) for the sauna and control groups.

With respect to the theta power, the repeated measures ANOVA indicated a significant main effect of set (F (3, 108) = 5.44, p < 0.01) and a significant interaction between group and set (F (3, 108) = 9.62, p < 0.01). We found no significant effects for electrode location or other interactions. A simple main effects test for set showed that the theta power significantly increased over time in the sauna group (F (3, 108) = 13.93, p < 0.01) but not in the control group (F (3, 108) = 1.14, n.s.). We further analyzed the simple main effect of set by conducting multiple comparisons using the Holm–Bonferroni method (MSe = 293.6901) to compare each set. The result showed a significant increase at post1, post2, and post3 compared with the pre-sauna phase (Table 1).

For the lower1 alpha power, the repeated measures ANOVA indicated a significant main effect of set (F (3, 108) = 7.67, p < 0.01), a marginally significant main effect of group (F (1, 36) = 3.31, p < 0.1), and a significant interaction between group and set (F (3, 108) = 8.10, p < 0.01). We found no significant effects for electrode location or other interactions. A simple main effects test for set showed that the lower1 alpha power significantly increased over time in the sauna group (F (3, 108) = 15.45, p < 0.01) but not in the control group (F (3, 108) = 0.32, n.s.). Using the Holm–Bonferroni method (MSe = 93.1421) to compare each set, we found a significant increase at post1, post2, and post3 compared with the pre-sauna phase (Table 1).

For the lower2 alpha power, the repeated measures ANOVA indicated a marginally significant main effect of set (F (3, 108) = 2.57, p < 0.1), a significant main effect of group (F (1, 36) = 5.21, p < 0.05), and a significant interaction between group and set (F (3, 108) = 5.18, p < 0.01). We found no significant effects for electrode location or other interactions. A simple main effects test for set showed that the lower2 alpha power significantly increased over time in the sauna group (F (3, 108) = 7.31, p < 0.01) but not in the control group (F (3, 108) = 0.43, n.s.). We further analyzed the simple main effect of set by conducting multiple comparisons using the Holm–Bonferroni method (MSe = 93.1421) to compare each set. The result showed a significant increase at post2 and post3 compared with the pre-sauna phase (Table 1).

For the upper alpha power, the repeated measures ANOVA indicated a significant main effect of set (F (3, 108) = 2.87, p < 0.05), a significant main effect of group (F (1, 36) = 6.33, p < 0.05), and a significant interaction between group and set (F (3, 108) = 5.85, p < 0.01). We found no significant effects for electrode location or other interactions. Further, a simple main effects test for set showed that the upper alpha power significantly increased over time in the sauna group (F (3, 108) = 8.24, p < 0.01) but not in the control group (F (3, 108) = 0.48, n.s.). We then used the Holm–Bonferroni method (MSe = 55.8500) to compare each set. The result showed a significant increase on the post2 and post3 compared with the pre-sauna phase (Table 1).

For the beta power, the repeated measures ANOVA indicated a significant main effect of group (F (1, 36) = 5.33, p < 0.05). We found no significant effects for other main effects or any interactions (Table 1).

### Effects of sauna on cognitive performance

To assess whether cognitive performance changed after sauna bathing, we calculated and compared the average RT for the oddball task in the pre-sauna and post-sauna phases for all participants in the sauna and control groups (Fig. 5A). The two-way repeated measures ANOVA indicated a marginally significant main effect of stage (F (1, 18) = 3.24, p < 0.1) and a significant marginally interaction between group and stage (F (1, 18) = 3.04, p < 0.1). A simple main effects test for the stage showed that the RT significantly decreased in the sauna group (F (1, 18) = 6.29, p < 0.05) but not in the control group (F (1, 18) = 0.00, n.s.).

**Fig. 5.**
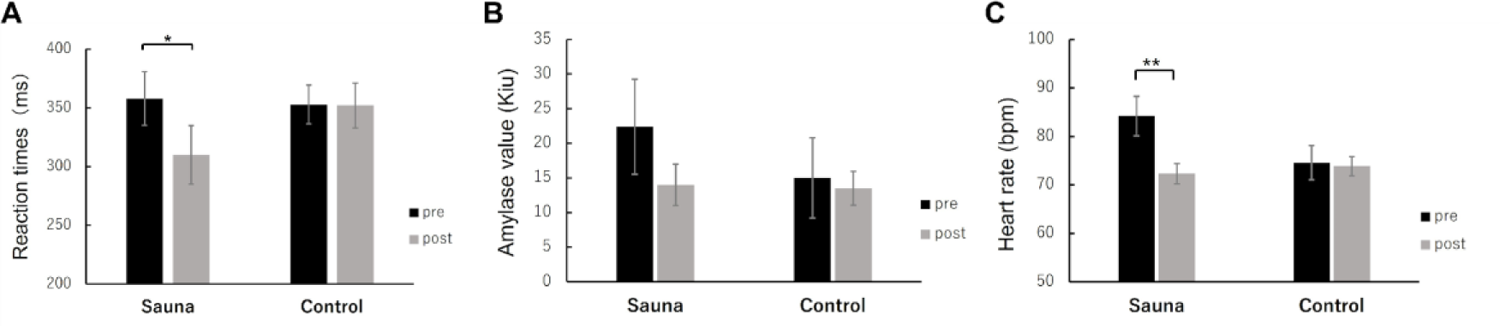
RT (A), Average SAA (B), and HR (C) before (pre) and after sauna (post3) in the sauna and control groups. Error bars represent s.e.m.* p < 0.05 and ** p < 0.01.

### Effects of sauna on physiological parameters

To assess whether physiological parameters changed after sauna bathing, we calculated and compared the SAA and HR in the pre-sauna and post-sauna phases for all participants in the sauna and control groups (Fig. 5B & 5C). The two-way repeated measures ANOVA for SAA indicated a marginally significant main effect of stage (F (1, 18) = 3.85, p < 0.1). We found no significant effects for group or interactions.

The two-way repeated measures ANOVA for HR indicated a significant main effect of stage (F (1, 18) = 24.13, p < 0.01) and a significant interaction between group and stage (F (1, 18) = 18.95, p < 0.01). A simple main effects test for the stage showed that the HR significantly decreased in the sauna group (F (1, 18) = 42.92, p < 0.01) but not for in control group (F (1, 18) = 0.16, n.s.).

### Effects of sauna on subjective feelings

To assess whether subjective feelings on participants’ physical and mental state changed by sauna bathing, we calculated and compared questionnaires’ scores in the pre-sauna phase and at each rest step in the sauna phase for all participants in the sauna and control groups (Figs. 6 and 7).

**Fig. 6.**
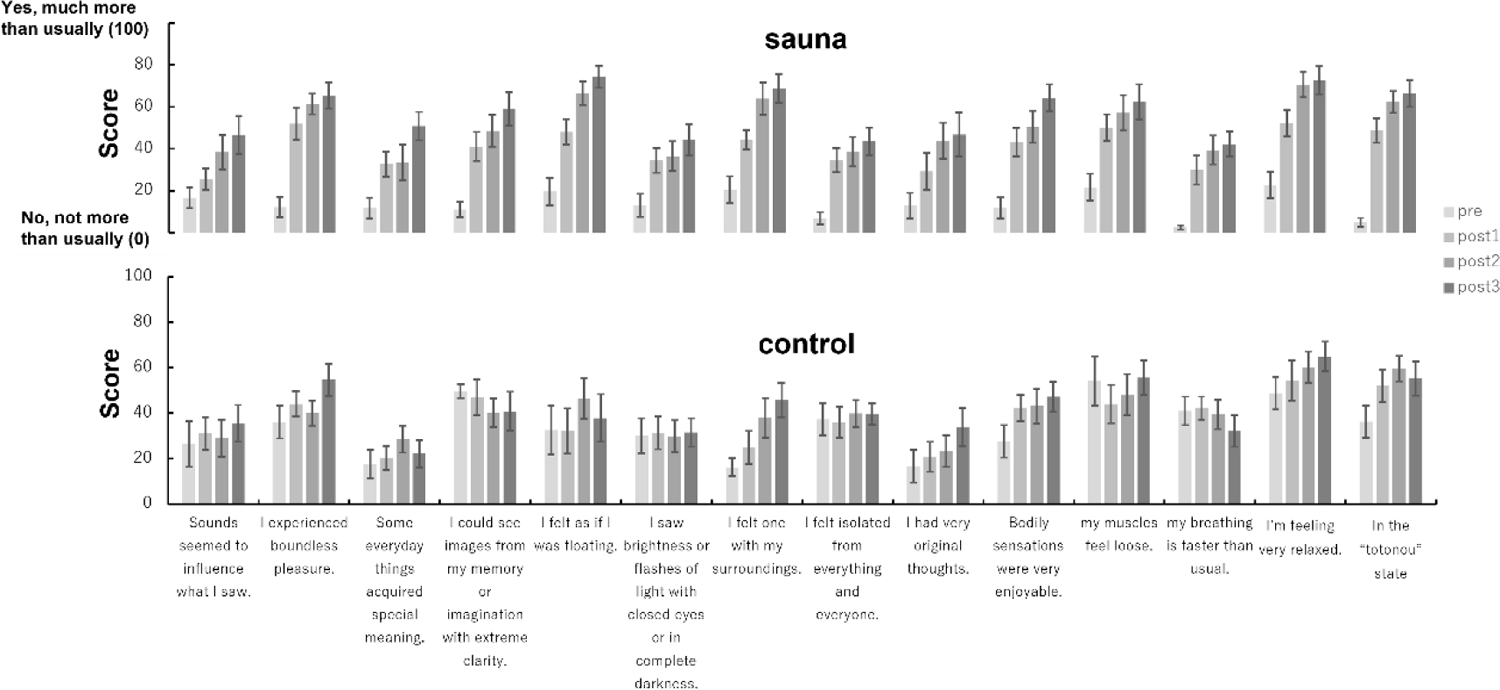
Average scores of questions in “Altered states of consciousness rating scale” before sauna (pre) and at each rest during the sauna set (post1, post2, post3) in the sauna and control groups. Error bars represent s.e.m.

**Fig. 7.**
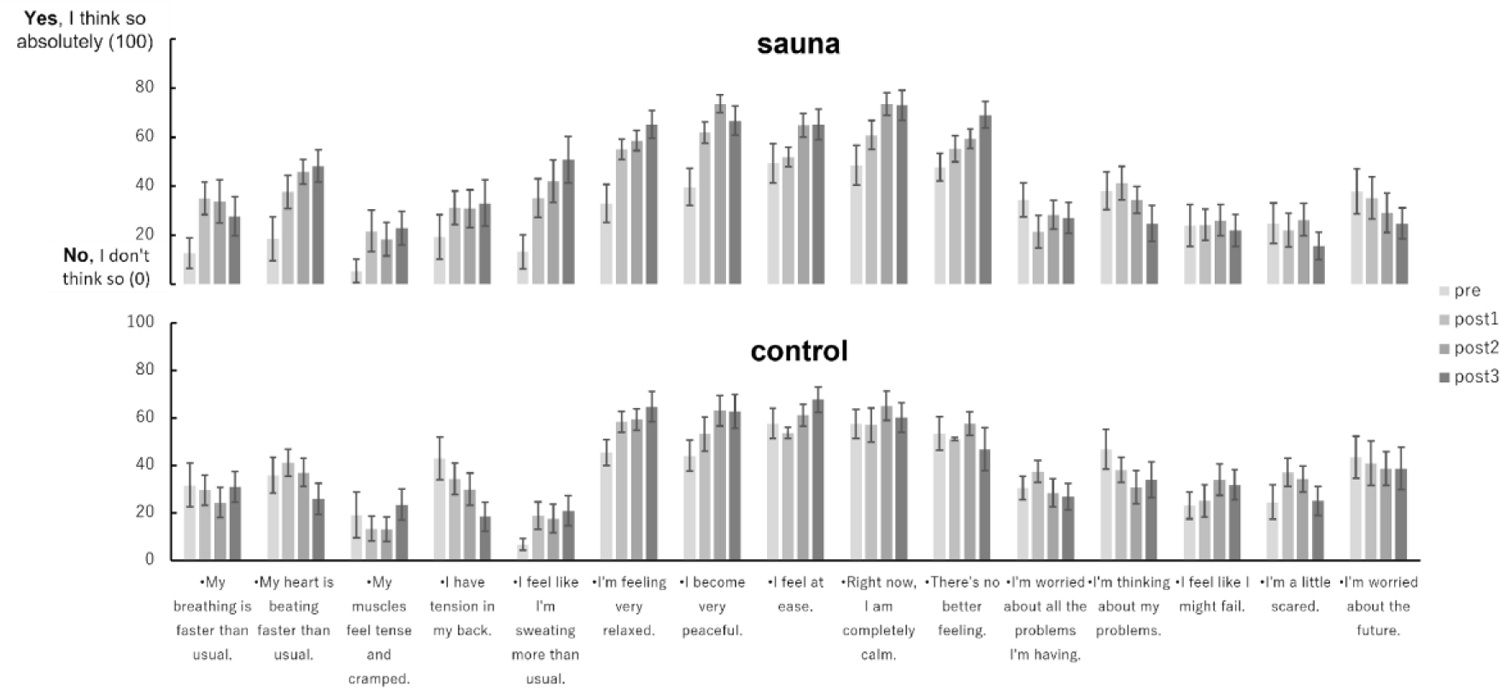
Average scores of questions in S-MARE before sauna (pre) and at each rest during the sauna set (post1, post2, post3) in the sauna and control groups. Error bars represent s.e.m.

The results of two-way repeated measures ANOVA are shown in Tables 2 and 3. In terms of subjective feelings, the altered states of consciousness rating scale and the S-MARE were used to assess whether subjective feelings on their physical and mental state changed through sauna bathing. In the altered states of consciousness rating scale, we observed significant changes in questions related to bodily sensations, such as *“Bodily sensations were very enjoyable*,” “*my muscles feel loose*,” “*I felt as if I was floating*,” etc. These statements indicate a relaxed body state. The answers to questions related to relaxation such as “*I’m feeling very relaxed*” also showed that people became more relaxed after sauna. Similarly, in another S-MARE questionnaire used to assess relaxation effects, the answers to the question “*There’s no better feeling*” showed a significant interaction between group and set, indicating that the sauna group was more relaxed after sauna; the responses to the two questions reflecting physiological tension physiological tension after the sauna, which might be related to the physiological reaction after the cold water. Significant main effects of set were observed in 4 of the 5 questions related to “psychological calm”. The most important was the answer to the question of being “*in the “totonou” state*,” which had a significant interaction and main effects of set. This indicated that the sauna group entered the “totonou” state after sauna, while the control group did not.

**Table 3.**
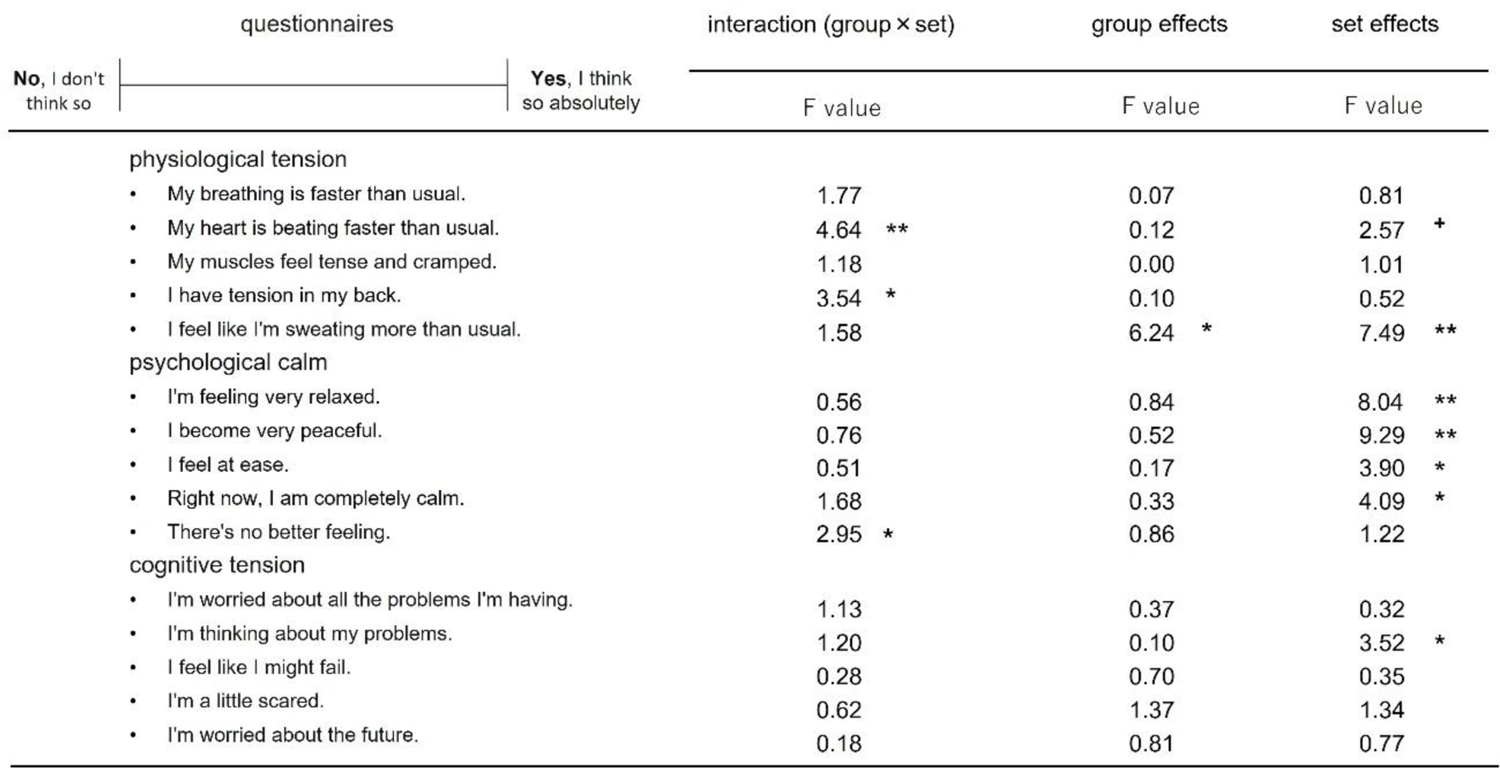
Results of two-way repeated measures ANOVA for all questions in S-MARE. **+** p < 0.1, * p < 0.05 and ** p < 0.01.

### AI decoding performance

We classified the data of all sets using our AI decoding algorithm. Figure 8 shows the classification accuracy of each participant in the sauna group; the average accuracy of the sauna group was 88.34% (SD = 6.71), which was significantly different from chance (50%), as tested by a one-sample t-test (t (9) = 18.08, p < 0.01).

**Fig. 8.**
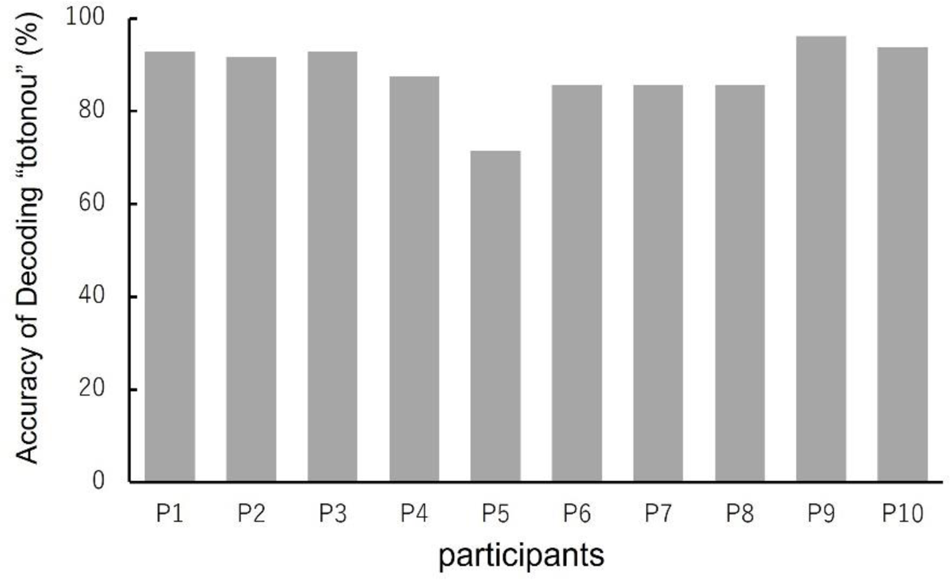
Classification accuracy of decoding “totonou” for each participant in the sauna group.

## Discussion

In the current study, to test the effects of the sauna bathing and evaluate the neural activity and body state in “totonou,” we conducted an experiment in which participants underwent three sets of alternating hot and cold baths in three steps: sauna (hot) → water bath (cold) → rest. First, the “totonou” state appeared at rest; thus, we measured the participants’ In-ear EEG during this period, finding a gradual increase in the theta and alpha power during the sauna phase. Second, we performed an auditory oddball task in pre-sauna and post-sauna phases to measure participants’ ERP using scalp EEG. The P300 amplitude as well as the reaction time decreased significantly after sauna, while that of MMN significantly increased. In addition, the participants’ subjective responses to the questionnaires showed significant changes in physical relaxation and other indicators over sauna bathing.

Our findings indicate that the P300 amplitude significantly decreased in the post-sauna phase compared to the post-sauna phase, which is consistent with the results of a previous study [14]. We also found that MMN amplitude increased significantly in the post-sauna phase compared with that in the pre-sauna phase. P300 is often linked with attention [17,18]. In fact, P300 amplitude has been suggested to reflect the amount of attentional resources allocated to the stimulus; therefore, the P300 is used as an indicator of the amount of attention allocated to a task [24,32]. Therefore, the decrease in P300 amplitude may be related to a decrease of attentional resources. On the other hand, the MMN is considered to reflect a pre-attentive auditory process [25,33]. In other words, MMN elicitation did not correlate with the attention as MMN can be elicited without the listener subjectively paying attention to the stimuli [21-23]. Previous studies have shown that MMN amplitude is closely related to auditory discrimination and can be used as an indicator of auditory discrimination accuracy [34,35]. In general, an increase in MMN amplitude represents increased sound discrimination. The sound stimulus used in the experiment was easily distinguishable to the participants; accordingly, there was no change in discrimination accuracy. However, the RT of participants in the sauna group who showed increased MMN amplitude in the post-sauna phase shortened compared to before the sauna. This suggests that participants became more sensitive to auditory stimuli throughout sauna bathing. This may lead to a decrease in P300 amplitude, since fewer attentional resources are needed for sound discrimination. These results indicate that sauna bathing may clear participants’ heads, making them more lucid. This is consistent with the results to the question *“I could see images from my memory or imagination with extreme clarity”* in the altered states of consciousness rating scale.

In terms of neural oscillation, we found that over each sauna set, the participants’ theta and alpha power gradually increased, while beta power did not change. Similarly, the alpha power was enhanced and the beta power unchanged in a previous study [14]; however, the theta power increased in our experiment while it did not change in their study. This difference may be due to differences in the experimental process because the EEG in the previous study was recorded at 90 min after the sauna while in ours In-ear EEG was measured during the sauna-phase. The results of the present study were very similar to a previous study on meditation [36], which demonstrated significant increases in theta and alpha power in the meditation condition compared to the rest condition. Higher theta activity probably reflects increased awareness and attention, as well as increased cognitive and emotional processing [37]. Another previous study on meditation showed that meditation reduced the HR level within participants more than rest [38]. In the present study, the sauna group also showed lower HR after sauna compared to the control group. This supports that the “totonou” state might be similar to that obtained after meditation. Moreover, these physiological results are consistent with those of questions related to emotional processing in the altered states of consciousness rating scale, such as “*I felt isolated from everything and everyone*,” “*I felt isolated from everything and everyone*,” and “*Some everyday things acquired special meaning*,” etc. On the other hand, the increased alpha activity may be related to relaxation [39,40]. This is consistent with the effects observed after high-intensity exercise, where an increase in alpha energy activity was accompanied by a high subjective experience of well-being and/or positive emotion [41]. This aspect was reflected in the questionnaire results as well. For example, the responses to questions *“I’m feeling very relaxed*,” “*I experienced boundless pleasure,”* etc. showing significant interactions between group and set.

We further subdivide alpha more into lower1 alpha, lower2 alpha, and upper alpha based on IAF. Lower1 alpha is associated with cognitive processes of internalized attention, alertness, and expectancy [42]. Changes in power of lower2 alpha and upper alpha are usually associated with tasks such as working memory or attention tasks. The lower-2 alpha band reflects expectancy whereas the upper alpha band is particularly relevant for processing semantic or task specific information [43,44]. In a previous study [45], the EEG power spectrum at rest was compared with that during memory tasks; during memory task there was a power reduction in the lower1 alpha, lower2 alpha, and upper alpha bands, indicating occupied brain resources. During the rest period in our experiment, participants did not participate in any task, they just closed their eyes and relaxed. The increase in power of lower1 alpha, lower2 alpha, and upper alpha bands observed during this period may indicate that brain resources are further released in the sauna-phase than in the pre-sauna phase.

From these results, we think that the “totonou” state is manifested in physical and mental feelings of relaxation, pleasure, mental clarity, accompanied by well-being and/or positive emotions. Changes in brain activity were reflected by increased theta and alpha power, decreased P300 and increased MMN amplitudes in the auditory discrimination task. The increase in MMN indicates increased activation of the pre-attentional auditory process, leading to a decrease in attention-related brain activity P300. This indicates that the brain is in a more efficient state.

Finally, the average performance of decoding the “totonou” state using the In-ear EEG reached 88.34%. This indicates the potential to classify the brain state using In-ear EEG and provides a reliable basis for future applications. For example, decoding-based neurofeedback could help users enter the “totonou” state not only in the sauna but also without sauna bathing. However, in this study, we did not conduct an experiment to achieve decoding-based neurofeedback. However, in our future studies, we will classify “totonou” and “no-totonou” states by AI decoding based on the present study, and conduct neurofeedback training on the user to verify whether the user can enter the “totonou” state. Accordingly, neurofeedback training will be conducted on the user to verify whether they can enter the “totonou” state without sauna bathing.

## Conclusion

In this study, the beneficial effects of sauna bathing were corroborated by both physiological data and a subjective emotional evaluation, which allowed us to characterize the “totonou” state as one of physical and mental relaxation, clarity of mind, and happiness and/or positive emotions. We further observed increased theta and alpha power as well as decreased P300 and increased MMN amplitudes. This manifests in objective physical changes (decreased HR) and cognitive responses (shorter RT). Finally, the average accuracy of classifying brain states using our AI decoding algorithm was 88.34%, having potential for future application.

## Acknowledgments

We express our sincere thanks to Mahiru Murakoshi, Karin Yamakawa and Daichi Inoue for helping to conduct the experiments. We would like to thank Enago (www.enago.jp) for the English language review.

